# gCoSRNA: Generalizable Coaxial-Stacking Prediction for RNA Junctions Using Secondary Structure

**DOI:** 10.1101/2025.07.26.666950

**Authors:** Shasha Li, Qianqian Xu, Ya-Lan Tan, Jian Jiang, Ya-Zhou Shi, Bengong Zhang

## Abstract

Coaxial stacking between adjacent stems is a key tertiary interaction that defines the spatial organization of RNA junctions, which are core structural motifs in folded RNAs. Accurate prediction of coaxial stacking is critical for RNA 3D structure modeling, yet existing computational tools remain limited, especially for junctions with variable numbers of branches or complex topologies. Here, we present gCoSRNA, a generalizable computational framework for predicting coaxial stacking configurations using RNA sequence and secondary structure as input. Instead of developing separate models for each junction type, gCoSRNA decomposes multi-way junctions into all possible adjacent stem pairs, termed pseudo two-way junctions, and uses a unified random forest classifier to evaluate stacking probabilities. Global stacking configurations are inferred by integrating these pairwise predictions, eliminating the need for explicit junction-type classification. Benchmarking on two independent test sets, including CASP15/16 and RNA-Puzzles targets, shows that gCoSRNA achieves consistently high accuracy (mean ∼0.87) across junctions with two to seven branches, outperforming existing junction-specific methods. These results highlight the model’s ability to capture higher-order structural features and its potential utility in RNA tertiary structure prediction pipelines. The source code is freely available at https://github.com/RNA-folding-lab/gCoSRNA.

## INTRODUCTION

RNA plays a central role in a wide range of biological processes, including gene regulation, catalysis, and molecular recognition, which critically depend on RNA’s ability to fold into complex three-dimensional structures (Morris and Mattick 2014; Mattick et al. 2023). Multi-way junctions, where three or more helices are connected through loop regions, are widespread in structured RNAs and play essential roles in their function (Lu et al. 2025; Serganov and Nudler 2013; Zhang 2024). For instance, hammerhead ribozymes adopt defined coaxial stacking patterns that bring helices into close proximity, creating compact active sites for catalytic self-cleavage (Lu et al. 2025). Riboswitches rely on similar multibranched folds to recognize small molecules and regulate downstream gene expression (Serganov and Nudler 2013). Transfer RNAs (tRNAs), with their conserved cloverleaf secondary structure, fold into a T-shaped three-dimensional conformation that enables accurate delivery of amino acids during translation (Zhang 2024). Understanding RNA structure is therefore not only essential for uncovering its biological functions, but also provides a foundation for RNA structure-based drug design and RNA nanotechnology (Yu et al. 2020; Guo 2010; Findeißet al. 2017).

Due to the high cost and limited scalability of experimental RNA structure determination (e.g., X-ray diffraction, nuclear magnetic resonance, and cryo-electron microscopy) (Zhang et al. 2022; Mukherjee et al. 2024; Li et al. 2025; von Löhneysen et al. 2024), some computational methods have been developed for predicting RNA 3D structures (Ou et al. 2022; Bu et al. 2025; Wang et al. 2023b; Zhang et al. 2024; Zeng et al. 2024; Wang et al. 2025). These approaches generally fall into three main categories: physics-based modeling (Watkins and Das 2023; Boniecki et al. 2016; Xia et al. 2013; Boudard et al. 2017; Shi et al. 2014, 2018; Wang et al. 2023c; Jin et al. 2019; Li and Chen 2023; Xiong et al. 2021; Tan et al. 2022; Zhang et al. 2021; Denesyuk and Thirumalai 2013), fragment assembly (Rother et al. 2011; Li et al. 2022; Xiong et al. 2023; Biesiada et al. 2016; Zhou et al. 2022), and AI–based methods (Wang et al. 2023a; Li et al. 2023; Sha et al. 2024; Shen et al. 2024; Kagaya et al. 2025; Li et al. 2018; Abramson et al. 2024). Although AI-based RNA structure prediction methods have advanced rapidly following the success of AlphaFold2 (Abramson et al. 2024; Jumper et al. 2021), recent results from CASP15 and CASP16 suggest that these approaches have not yet outperformed traditional physics-based and fragment assembly methods (Das et al. 2023; Kretsch et al. 2025). CASP16 further emphasized that RNA 3D structure prediction remains largely template-dependent (Kretsch et al. 2025). However, the number of experimentally determined RNA structures, particularly those containing multi-way junctions, remains limited, constraining the accuracy of fragment-based methods for complex RNAs (Zhang et al. 2024; Zeng et al. 2024). While coaxial stacking between stems connected through junctions is a key interaction shaping RNA topology, accurately modeling this interaction from physics remains a significant challenge (Butcher and Pyle 2011; Tinoco and Bustamante 1999; Laing et al. 2009). This highlights the need for reliable computational tools specifically designed to predict coaxial stacking in RNA junctions, which could be further integrated into physics-based frameworks to improve the overall accuracy of RNA 3D structure prediction (Zhang et al. 2022; Mukherjee et al. 2024).

To classify RNA three-way junctions, Lescoute and Westhof proposed an algorithm that groups junctions into three families based on sequence and base-pairing patterns, derived from 32 known RNA structures (Lescoute and Westhof 2006). Similarly, Laing and Schlick analyzed 62 RNAs with four-way junctions and identified nine structural families, outlining key features of each (Laing and Schlick 2009). Junctions within the same family share a common coaxial stacking arrangement, allowing the coaxial stacking of unknown RNAs to be inferred through family-level classification (Laing et al. 2012; Lamiable et al. 2012; Ali et al. 2023). However, the predictive power of these methods is limited by the small number of available training structures and the relatively coarse nature of family-level classification (Bu et al. 2025; Wang et al. 2023; Zhang et al. 2024). To directly predict coaxial stacking from sequence and secondary structure, Laing et al. developed Junction-Explorer, which combines a set of secondary structure features with a random forest classifier to predict coaxial stacking in three- or four-way junctions (Laing et al. 2012). However, the method requires separate models for different junction types, making it highly dependent on the availability of structurally resolved homologs (Laing et al. 2012).

In this study, we present gCoSRNA, a generalizable machine learning framework for predicting coaxial stacking in RNA junctions based solely on secondary structure and sequence information. Unlike previous approaches that rely on junction-specific models or family-level classifications, gCoSRNA decomposes each multi-way junction into all possible stem–stem pairs (defined as pseudo two-way junctions, e.g., H1–H2, H2–H3, H3–H1 in a three-way junction with three stems of H1, H2 and H3). A single unified model is trained to estimate stacking probabilities for individual stem pairs, which are then integrated to infer the global coaxial stacking configuration of any multi-way junction. This pairwise decomposition strategy effectively expands the training data and enables generalization across diverse junction types. By leveraging readily available secondary structure features, gCoSRNA provides a robust and scalable solution for coaxial stacking prediction, with potential applications in RNA 3D structure modeling and the study of RNA architectural organization.

## RESULTS

### Overview of gCoSRNA

gCoSRNA is a machine learning-based framework for predicting coaxial stacking patterns in RNA multi-way junctions using secondary structure-derived features. As illustrated in Fig. 1A, the framework consists of two main stages: (1) each multi-way junction is decomposed into pseudo two-way junctions formed by adjacent stem pairs (e.g., three pairs for a three-way junction, see Fig. S1 in the Supplementary Materials), and stacking probabilities are predicted for each pair using a pre-trained Random Forest (RF) model; (2) the pairwise predictions are then integrated to reconstruct the complete coaxial stacking configuration of the junction. The RF model was trained on experimentally resolved RNA 3D structures using features extracted from both primary sequence and secondary structure (Fig. 1B), including loop lengths and thermodynamic stability (Fig. 1C). Model parameters were optimized via tenfold cross-validation to ensure robustness and generalizability. This decomposition-based strategy enables gCoSRNA to achieve accurate and scalable predictions across a wide range of RNA junction architectures. Further methodological details are provided in the Materials and Methods section.

**Figure 1.**
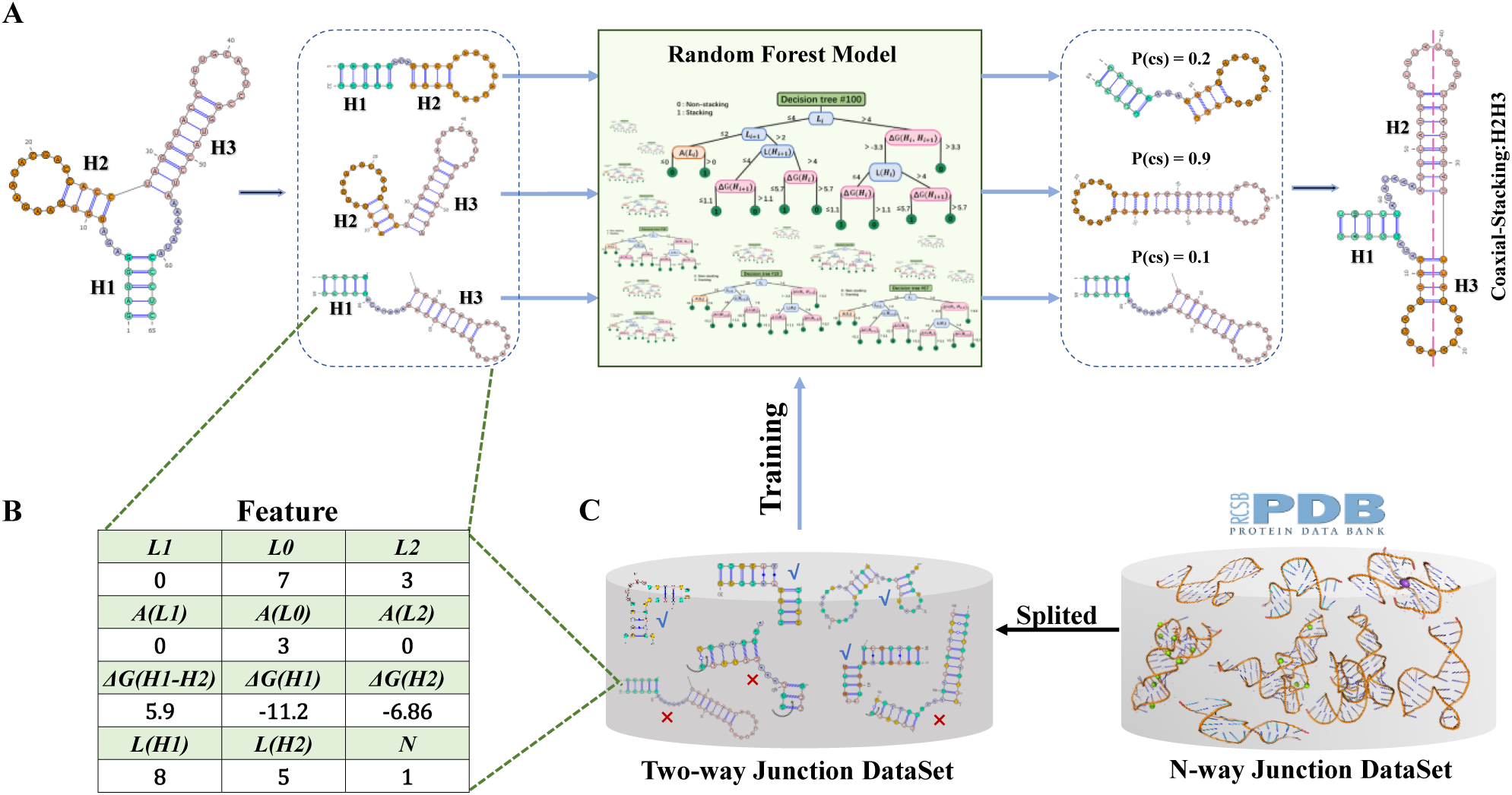
Overview of the gCoSRNA framework for coaxial stacking prediction. **A.** Schematic workflow of gCoSRNA. A representative RNA three-way junction with three helices (H1, H2, H3) is decomposed into three adjacent helix pairs (H1–H2, H2–H3, H1–H3) based on secondary structure. Each pair is independently evaluated by a pre-trained Random Forest (RF) model to estimate the probability of coaxial stacking. These pairwise probabilities are then integrated to infer the overall stacking configuration of the junction (e.g., coaxial stacking between H2 and H3). **B.** Example feature set used for each helix pair, incorporating sequence- and secondary structure-based features. **C.** Training pipeline of the RF model using features derived from experimentally resolved RNA 3D structures.

### Reproducibility of Junction-Explorer

RNA Junction-Explorer is a well-regarded method for predicting coaxial stacking in RNA multi-way junctions (Laing et al. 2012). However, as its original web server is currently unavailable, we re-implemented the model to enable direct comparison with gCoSRNA (see Materials and Methods for implementation details). To validate our reimplementation, we retrained Junction-Explorer using the original dataset, feature set, and hyperparameters described in Laing et al. (2012). Tenfold cross-validation yielded accuracies of 81% for three-way and 77% for four-way junctions, closely matching the published results (Laing et al. 2012) and confirming the fidelity of our implementation.

We further trained an enhanced version, Junction-Explorer-n, using our newly curated and expanded junction dataset. To ensure fair comparison on the original dataset, we conducted 10 rounds of random subsampling, each time withholding one-tenth of the original samples as a test set. The average prediction accuracies for three-way and four-way coaxial stacking reached 82% and 79%, respectively, showing improvements of approximately ∼1.2% and ∼2.6% over the original model. These results suggest that training with more comprehensive data can improve Junction-Explorer’s performance, though the gains are relatively modest (i.e., <3%).

### gCoSRNA performance evaluation

We first assessed the predictive performance of gCoSRNA using 10-fold cross-validation on the training set of pseudo two-way junctions. As shown in Fig. S2 in the Supplementary Materials, the model achieved consistently high accuracy across all folds, with an average area under the ROC curve (AUC) of 0.97 and fold-wise AUCs exceeding 0.96. Other evaluation metrics, including accuracy, precision, recall, F1-score, and Cohen’s Kappa, also demonstrated strong and stable performance, e.g., all folds yielded accuracy and F1-scores above 0.90, with recall consistently above 0.95 and precision near 0.90. These results highlight the strong discriminative power and stability of gCoSRNA in identifying coaxial stacking from pseudo two-way junction data.

Although gCoSRNA was trained exclusively on pseudo two-way junctions, it can be readily applied to RNA junctions of arbitrary order by predicting the coaxial stacking status of each pair of adjacent stems. Two-way junctions have a binary outcome, stacked or unstacked. For three-way junctions, four possible configurations exist: H1–H2, H2–H3, H3–H1, or no stacking (Fig. S3). Four-way junctions exhibit seven distinct patterns, including four single-stack cases (H1–H2, H2–H3, H3–H4, H4–H1), two double-stack cases (H1–H2 & H3–H4, H2–H3 & H4–H1), and a no-stacking configuration. For these junctions (order < 5), a prediction was considered correct only if the full coaxial stacking pattern matched the known annotation exactly. For junctions with five or more stems, where the number of possible configurations grows rapidly, we focused instead on pairwise accuracy (PA); see Eq. 3 in the Materials and Methods section. This metric assesses whether the stacking prediction for each adjacent stem pair is correct, providing a tractable means of evaluating performance in high-order junctions. We evaluated the predictive performance of gCoSRNA on the two independent test sets (Table 1).

**Table 1.** Prediction accuracy of different methods on two benchmark test sets.

| Test sets <sup>a</sup> |  | Size | Junction-Explorer | Junction-Explorer-n | gCoSRNA |
| --- | --- | --- | --- | --- | --- |
| Testset I | Two-way | 220 | - <sup>b</sup> | - | 0.94 |
|  | Three-way | 50 | 0.76 | 0.79 | 0.88 |
|  | Four-way | 40 | 0.67 | 0.70 | 0.85 |
|  | Five-way | 13 | - | - | 0.92 |
|  | Six-way | 45 | - | - | 0.97 |
|  | Seven-way | 49 | - | - | 0.95 |
| Testset II | Two-way | 80 | - | - | 0.86 |
|  | Three-way | 9 | 0.33 | 0.44 | 0.56 |
|  | Four-way | 29 | 0.58 | 0.77 | 0.86 |

<sup>a</sup> For two- to four-way junctions, the reported values represent full-configuration prediction accuracy. For junctions beyond five-way level, pairwise accuracy (PA) is used. <sup>b</sup>“-” indicates that the prediction was not performed due to the absence of a corresponding trained model/parameters for the specified junction type in the original RNA Junction-Explorer implementation

#### On Testset I

Testset I comprised a diverse collection of RNA junctions with a distribution similar to the training set (see Training and Test datasets in the Materials and Methods). As shown in Table 1 and Table S1 in the Supplementary Material, gCoSRNA achieved its highest performance on two-way junctions, with an AUC of 0.91 (Fig. 2A) and an overall accuracy of 0.94 (F1-score of 0.96; see Table S1), highlighting the consistency between native and pseudo two-way stacking features. The confusion matrix reveals that gCoSRNA correctly predicted 203 out of 206 coaxially stacked helix pairs (Fig. 2A). However, only one-third of non-stacked pairs were accurately classified, likely due to the strong class imbalance in the test set, where stacked configurations were heavily overrepresented relative to non-stacked ones (Fig. S4).

**Figure 2.**
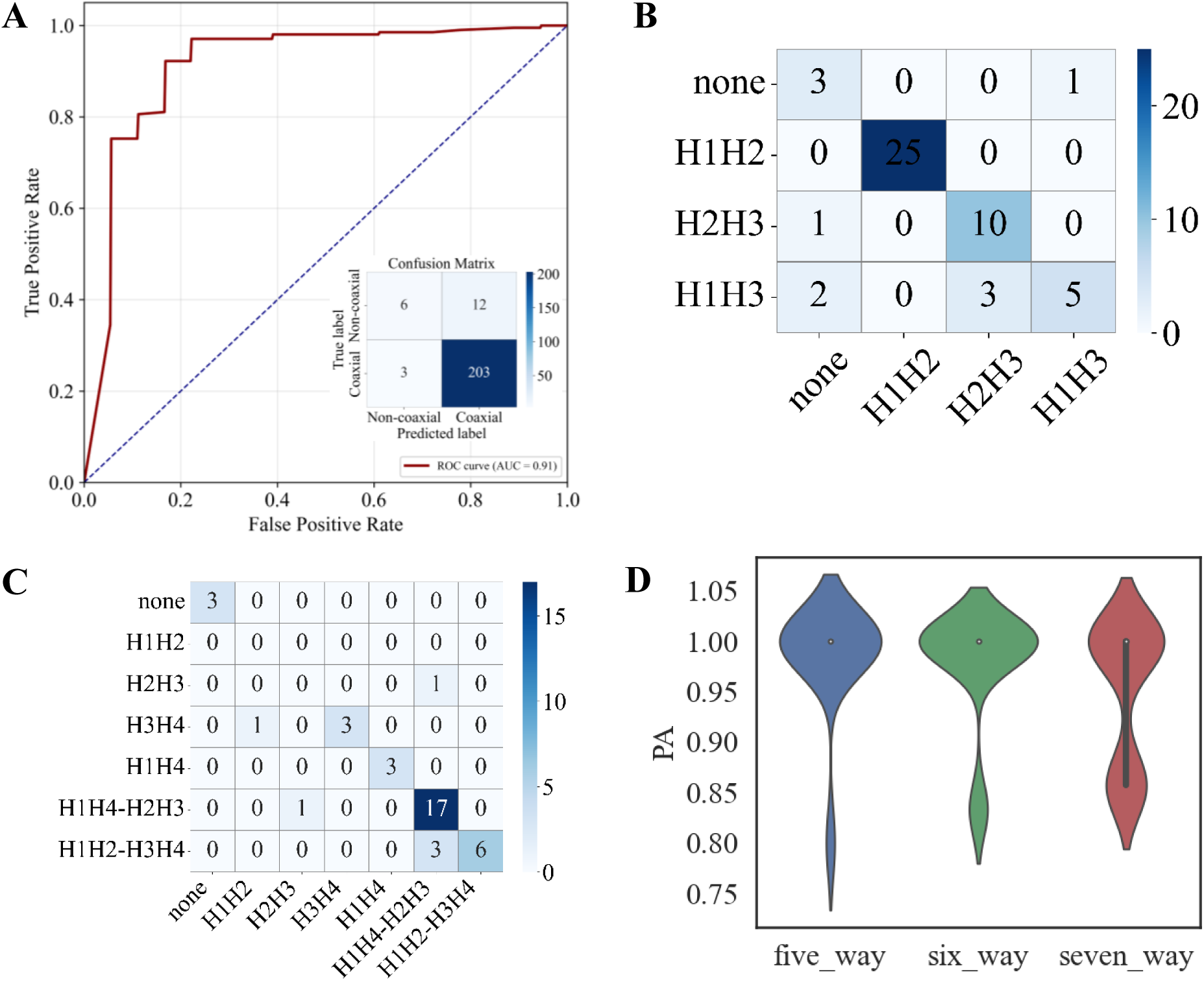
Performance of gCoSRNA on Test Set I. **A.** ROC curve for coaxial stacking prediction in two-way junctions, with the corresponding confusion matrix shown as an inset. **B&C.** Confusion matrices for coaxial stacking prediction in three-way (B) and four-way (C) junctions. **D.** Distribution of pairwise accuracy (PA) values for five-, six-, and seven-way junctions, visualized as violin plots.

For three-way and four-way junctions, gCoSRNA achieved overall stacking configuration accuracies of 0.88 and 0.85, respectively (Table 1), along with consistently high average F1-scores in the pairwise pseudo two-way classification tasks (Table S1). Confusion matrix analyses (Figs. 2B and 2C) revealed that most stacking configurations were accurately predicted, although certain arrangements, such as H1–H3 in three-way and H1H2–H3H4 in four-way junctions, showed reduced sensitivity, likely due to local structural ambiguity or limited representation in the training data. These findings support the effectiveness of decomposing junctions into adjacent stem pairs for accurate reconstruction of global coaxial stacking topologies.

For five-way junctions, gCoSRNA achieved a pairwise accuracy (PA) of 0.92 and an F1-score of 0.89. Notably, despite the limited number of RNA-only higher-order junctions in the training set, the model maintained high predictive accuracy on six- and seven-way junctions in the test set (n = 94). After removing redundancy with the training data and among test samples, gCoSRNA attained an average PA of 0.95 across these complex junctions (Table 1; Fig. 2D). Representative examples of five-, six-, and seven-way junctions with distinct coaxial stacking configurations are shown in Fig. S5, where gCoSRNA accurately recovered nearly all native stacking interactions, with only minimal false positives. These results highlight the model’s strong extrapolation capacity to structurally diverse and underrepresented junction types, supporting its broad utility in RNA tertiary structure modeling.

#### On Testset II

Test Set II was curated from RNA-Puzzles and CASP RNA targets, which are structurally diverse RNAs with no significant sequence overlap with the training set. Among the 80 two-way junctions, gCoSRNA achieved 0.86 accuracy (Table 1) and 0.90 F1-score (Table S1), indicating strong generalizability to unseen RNA structures. Performance on 9 three-way junctions dropped to 0.56 accuracy and 0.57 F1-score, likely due to limited sample size and increased structural variability, particularly within RNA–ligand/ion interactions (see Fig. S6). In contrast, prediction on 29 four-way junctions remained robust, with 0.86 accuracy (Table 1), underscoring the model’s ability to handle moderate structural complexity. These results highlight both the adaptability and current limitations of gCoSRNA, suggesting that incorporating RNA–protein complex data during training may further improve generalization in future applications.

### Comparison with existing methods

To further evaluate the performance of gCoSRNA, we compared it against two versions of Junction-Explorer. Both models were applied to the three-way and four-way junctions in Testset I and Testset II (Table 1). Consistent with the training set results, Junction-Explorer-n achieved slightly better average accuracy (0.71) than the original Junction-Explorer (0.64) three- and four-way junctions, but both were significantly worse than gCoSRNA (0.79). In Testset I, gCoSRNA attained accuracies of 0.88 and 0.85 on three-way and four-way junctions, respectively. Compared to Junction-Explorer-n, these represent improvements of 11% and 21%, and relative to Junction-Explorer, the gains were 16% and 27%, respectively. In Testset II, although gCoSRNA’s accuracy on three-way junctions (0.56) was lower than that in Testset I (0.88), it still notably outperformed both Junction-Explorer (0.33) and Junction-Explorer-n (0.44). Specifically, among the 9 three-way junctions, gCoSRNA correctly predicted 5, while Junction-Explorer-n correctly predicted only 4. There were 3 RNAs for which both methods failed, possibly due to ligands/ion-induced conformational shifts or tertiary constraints not captured by the training data (Fig. S6). For the 29 four-way junctions, gCoSRNA and Junction-Explorer-n achieved correct predictions on 25 and 22 cases, respectively. Overall, these comparisons reveal that building junction-specific models (e.g., for three-way or four-way junctions) does not confer a clear advantage over gCoSRNA, which reconstructs the global stacking configuration by independently predicting pairwise stem-stackings. This suggests that the abundant and structurally diverse two-way junctions (native or pseudo) can serve as effective training surrogates for learning generalizable coaxial stacking rules.

### Feature Importance Analysis

To investigate the contribution of individual features to gCoSRNA’s performance, we analyzed feature importance scores derived from the trained RF model using two standard split-based criteria: Gini impurity and Information Gain; see Fig. S7 in the Supplementary Materials. Both metrics evaluate how effectively a feature partitions the data, with higher scores indicating greater contributions to the model’s decision-making process across the ensemble. As shown in Fig. S7, the most important features are the coaxial stacking free energy between adjacent stems (*ΔG(H1-H2*)), the loop length connecting these stems (*L0*), the number of intervening stems between them in the original n-way junction (*N*), and the free energy of the first stem (*G(H1)*). These results align with established biophysical principles that stacking interactions are primarily determined by thermodynamic stability and relative spatial arrangement within the junction (Laing et al. 2012; Mathews et al. 2004).

## DISCUSSIONS

The design of gCoSRNA is grounded in the principle of structural modularity, wherein complex multi-branch junctions are decomposed into pseudo two-way junctions units. This strategy offers two main advantages. First, it obviates the need to train separate models for each junction order, enabling a single unified binary classifier to be applied across diverse junction types. Second, it substantially expands the effective training dataset by extracting numerous pseudo two-way units from higher-order junctions, addressing the common challenge of data scarcity in multi-way junction modeling. These factors contribute to the model’s strong robustness and generalizability across a wide range of RNA structures.

To further assess the validity of this decomposition-reconstruction strategy, we conducted a case study based on feature similarity analysis. Specifically, we calculated a cosine similarity score (*Fs*) between the 12-dimensional feature vectors of pseudo two-way junctions in the test set and all samples in the training set. As illustrated in Fig. 3, we examined a representative example from the test set, a four-way junction (8t2p_1) containing two native coaxial stacking interactions: H1–H4 and H2–H3. We extracted two pseudo two-way motifs (H1–H2 and H3–H4) and compared each to the training set. The five most similar samples for each pseudo motif were visualized in 2D and 3D. Strikingly, all top matches originated from real two-way junctions (i.e., hairpins with internal loops or bulges) rather than from other four-way junctions. This finding demonstrates that key stacking features are shared between pseudo junctions derived from complex multibranch loops and native two-way junctions. It highlights that gCoSRNA effectively learns generalizable stacking principles that transfer across different topologies.

**Figure 3.**
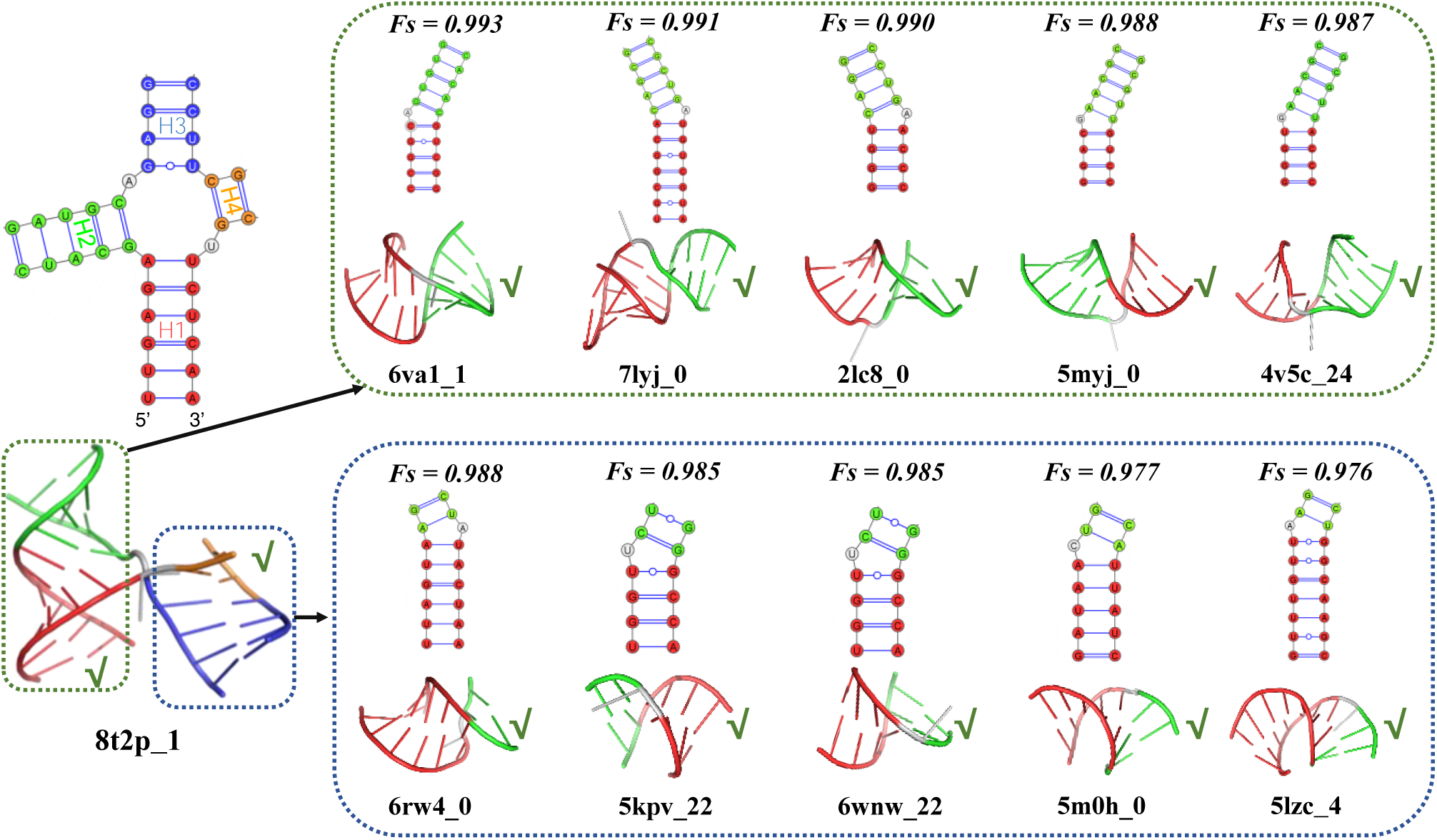
Positive impact of the modular decomposition strategy used in gCoSRNA. A &. **B.** 2D and 3D structures of the top five training-set junctions most similar (based on feature vectors) to the H1–H4 (top) and H2-H3 (bottom) pseudo two-way junctions derived from the four-way junction 8t2p_1. In all 2D and 3D visualizations, stem elements are color-coded consistently to highlight structural correspondence.

Despite the overall effectiveness of the decomposition-based strategy, certain mispredictions highlight its limitations. As shown in Fig. 4, the RNA 4wfm_1 contains a three-way junction in which H1–H3 forms a coaxial stacking interaction in the native structure, yet gCoSRNA incorrectly predicts stacking between H1–H2. To understand this error, we performed a feature similarity analysis. For the H1–H2 pair, all five closest training instances (four from canonical two-way junctions and one from a four-way junction) exhibit coaxial stacking, likely biasing the model’s decision. In contrast, among the top five matches for H1–H3, only two show coaxial stacking, and the remaining three (from four-way junctions) do not. These suggest that the model’s learned decision boundaries may be skewed when the pseudo two-way motifs originate from topologically distinct contexts, highlighting a trade-off between generalizability and structural specificity in the decomposition-based prediction framework.

**Figure 4.**
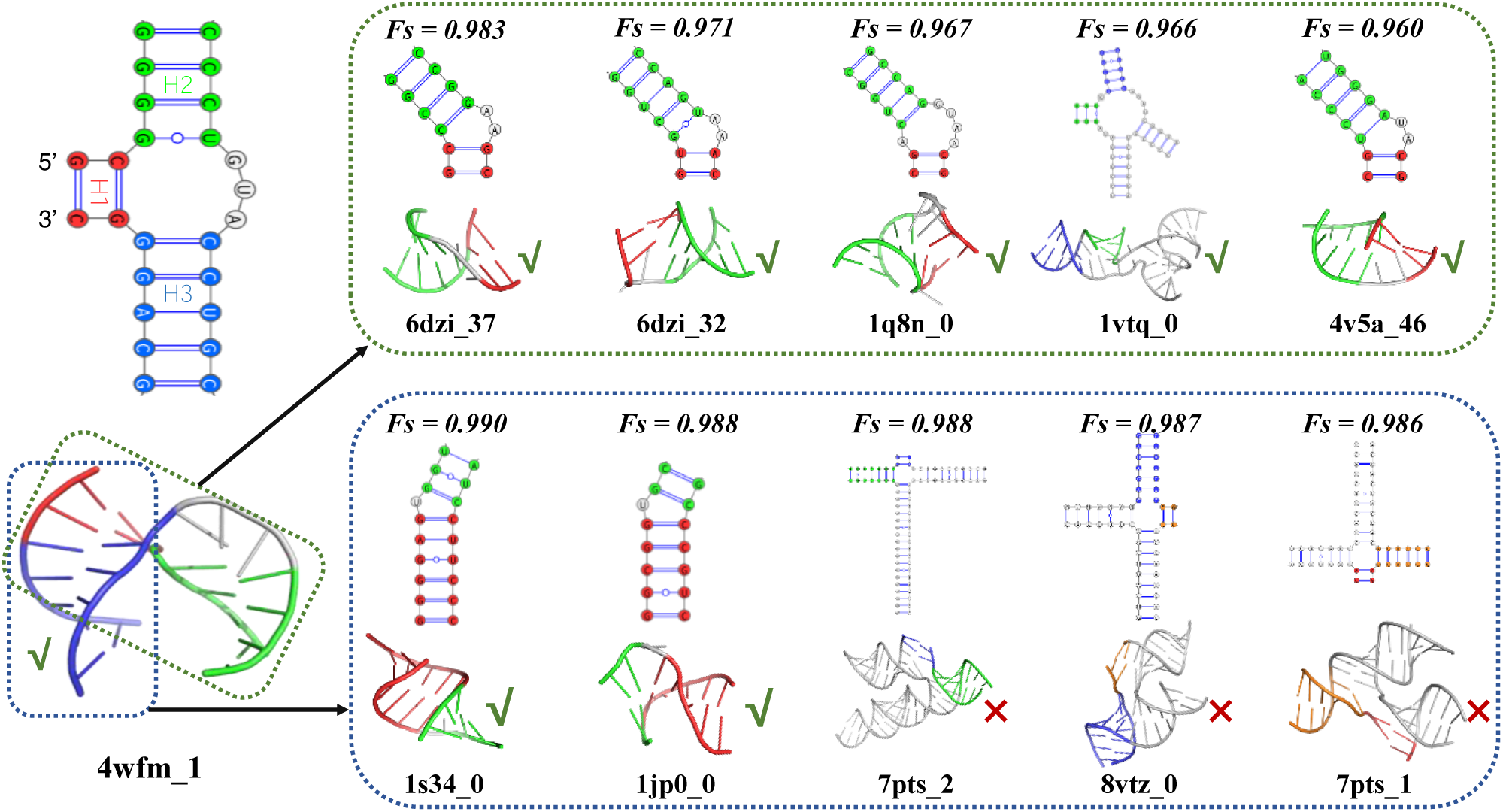
Limitations of the modular decomposition strategy in gCoSRNA. 2D and 3D structures of the top five training-set junctions most similar to the H1–H2 (top, red & green) and H1–H3 (bottom, red & blue) pseudo two-way junctions derived from the three-way junction 4wfm_1. All stem elements are consistently colored across 2D and 3D representations.

Given that coaxial stacking plays a critical role in shaping the 3D topology of RNA junctions, accurate identification of stacking interactions is expected to enhance structural modeling accuracy (Wang et al. 2023; Tinoco and Bustamante 1999). For instance, in our previous work, we developed a sequence-based coarse-grained model for RNA 3D structure prediction (Shi et al. 2018; Wang et al. 2023; Shi et al. 2014), and found that incorporating coaxial stacking interactions significantly improved both structural accuracy and thermodynamic stability, particularly for two-way junctions (Jin et al. 2019; Shi et al. 2015; Zhang et al. 2019). To further evaluate the contribution of coaxial stacking prediction in more complex RNA structures, we conducted a comprehensive analysis on three- and four-way junctions from Test set II, which includes structure prediction models submitted to CASP15/16 and RNA-Puzzles. As shown in Fig. 5A (left), the overall distribution of RMSD values between predicted and native structures spans a wide range (∼4–50 Å), with a mean of ∼27.0 Å, reflecting large variability in structural accuracy. To control for differences in secondary structure, we first extracted base-pairing patterns using DSSR and selected only those models with a secondary structure F1-score > 0.8. These models displayed a more compact RMSD distribution (Fig. 5A, middle), with a reduced mean RMSD of ∼17 Å, representing a 59% improvement over the full set. From this secondary-structure-consistent subset, we further identified models that correctly reproduced the coaxial stacking configurations predicted by gCoSRNA. This final group exhibited the most concentrated RMSD distribution (Fig. 5A, right), with a mean RMSD of ∼10 Å and the majority of models falling below 15 Å, yielding a ∼70% improvement relative to all models. As illustrated in Fig. 5B–D, for three representative multi-way junctions, incorporating coaxial stacking information led to a substantial improvement in structural accuracy, reducing the RMSD from over 14 Å to approximately 5 Å. It is worth noting, however, that the RNAs used for benchmarking often contain additional structural elements such as hairpins or single-stranded regions beyond the junction itself. Moreover, the global topology of a multi-way junction may be influenced not only by the stems involved in coaxial stacking, but also by the angular relationships between them and other flanking stems within the junction (see Fig. S8). As a result, the incorporation of stacking information does not always lead to a comparable reduction in RMSD across all RNA targets, as observed in the representative examples. Nevertheless, these results underscore the importance of coaxial stacking as a higher-order structural determinant and demonstrate that gCoSRNA’s stacking predictions can serve as a reliable criterion for selecting high-quality 3D models from structure prediction pipelines. Therefore, integrating gCoSRNA into existing RNA structure prediction frameworks, either as a constraint in potential energy functions or as a post hoc model evaluation method, holds strong potential to predict RNA 3D structures with high accuracy.

**Figure 5.**
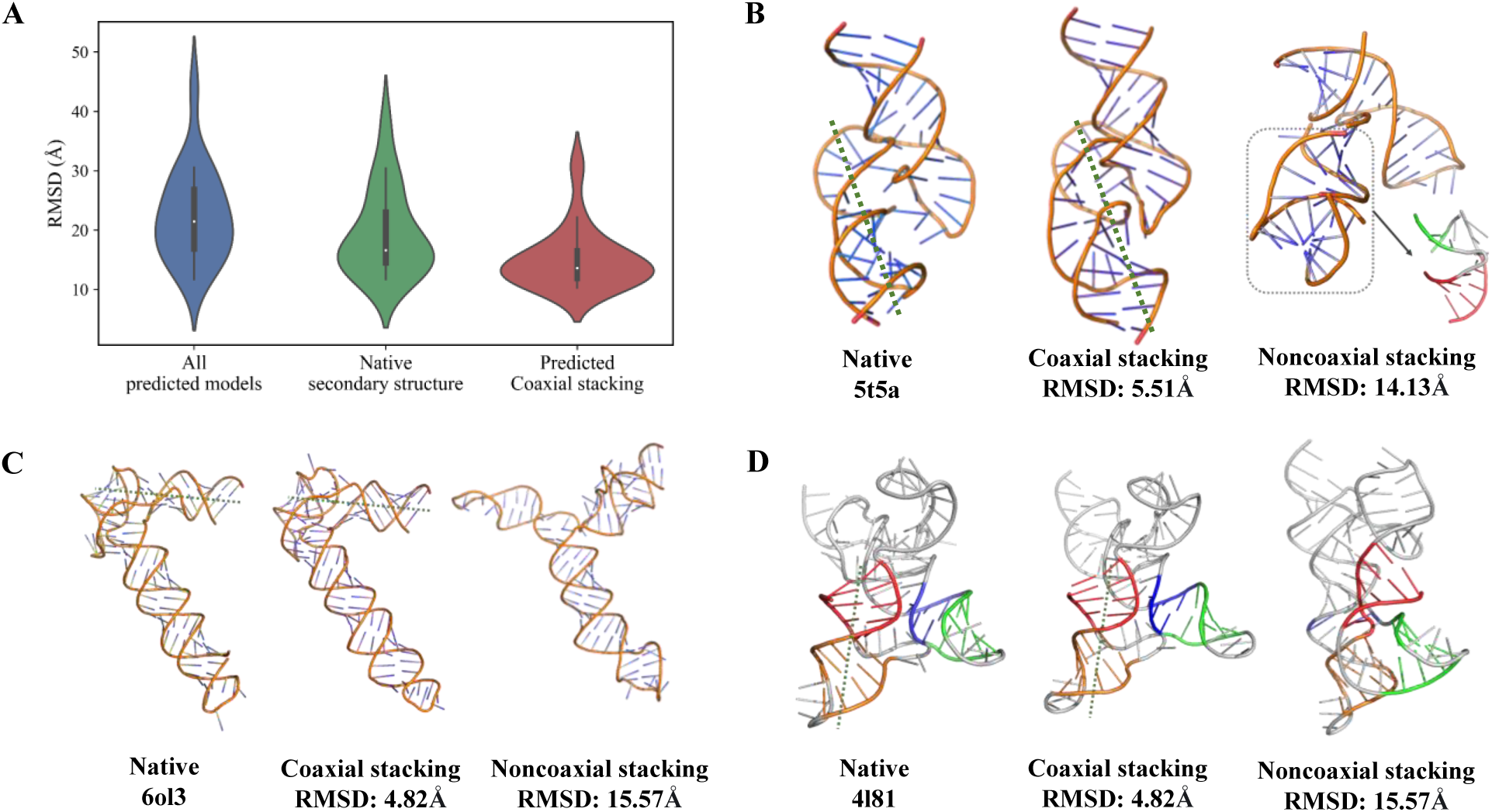
Coaxial stacking prediction as an indicator of 3D model accuracy. **A.** RMSD distributions of predicted RNA 3D structures for three- and four-way junctions in Test Set II, which includes CASP15/16 and RNA-Puzzles targets. From left to right: all submitted models (blue), models with high secondary structure fidelity (F1 score >0.8, green), and models that additionally match the coaxial stacking configuration predicted by gCoSRNA (red). **B–D.** Representative examples of three junctions, each showing three structures (from left to right): the native structure, a predicted model with coaxial stacking consistent with gCoSRNA predictions, and a model with inconsistent stacking.

## CONCLUSION

In this study, we developed gCoSRNA, a generalizable and topology-independent framework for predicting coaxial helical stacking in RNA multi-way junctions using only secondary structure-derived features. By decomposing complex junctions into pseudo two-way stem pairs and applying a unified machine learning model, gCoSRNA captures shared stacking signatures across diverse junction types and achieves accurate predictions without relying on junction-specific classifications. Systematic evaluations on 3- to 7-way junctions demonstrate strong predictive performance and robust generalization. When integrated into 3D modeling workflows, gCoSRNA’s stacking predictions substantially improve structural fidelity, highlighting the biological relevance of coaxial stacking as a key determinant of RNA tertiary topology.

Despite its strong performance, several limitations remain. First, although structurally diverse two-way junctions, such as hairpins containing bulges or internal loops, contribute to improved generalization, coaxial stacking cases in the PDB outnumber non-stacking instances by approximately sixfold (Fig. S10). This substantial class imbalance may predispose the model to favor stacking predictions (Fig. 4). Second, due to the limited availability of non-redundant RNA 3D structures, gCoSRNA (similar to Junction-Explorer (Laing et al. 2012)) relies on manually crafted features derived from secondary structure. While such features offer interpretability (e.g., coaxial stacking free energy and connected loop length between two stems are identified as key contributors; see Fig. S7), they may constrain predictive power and scalability. With future increases in available data, adopting deep learning strategies for automated feature extraction may enhance prediction accuracy and enable sequence-level coaxial stacking inference (Shen et al. 2024; Abramson et al. 2024; Wang et al. 2025; Chaturvedi et al. 2025; Chang et al. 2024). Finally, although benchmarking on CASP and RNA-Puzzles targets shows that gCoSRNA can help identify structurally accurate models, the optimal strategy for integrating coaxial stacking predictions into existing RNA folding frameworks, particularly physics-based or deep learning–driven methods, remains an open question (Mukherjee et al. 2024; Xu and Chen 2020; Mustoe et al. 2012; Mustoe et al. 2014). Addressing these challenges will be essential to further advancing RNA tertiary structure prediction and functional inference.

## MATERIALS AND METHODS

### Pairwise Decomposition of Multi-way Junctions

In contrast to RNA Junction-Explorer, which treats the entire secondary structure of each RNA junction as a single training sample (Laing et al. 2012), gCoSRNA decomposes every *n*-way junction into multiple pseudo two-way junctions, each serving as an independent sample. Specifically, a pseudo two-way junction is defined as a pair of adjacent stems along with all intervening unpaired nucleotides (e.g., loop regions) that connect them. As illustrated in Fig. S1, a three-way junction with stems H1, H2, and H3 is decomposed into three pseudo two-way junctions: H1–H2, H2–H3, and H3–H1. Each pseudo two-way junction is assigned a binary label indicating whether its two stems are coaxially stacked in the native 3D structure.

### Feature Extraction from Pseudo Two-way Junctions

For each pseudo two-way junction (comprising stems H1 and H2), we extracted 12 features derived from sequence and secondary structure information (Laing et al. 2012):

1. *L0*: Length of the loop directly connecting H1 and H2. For true two-way junctions (i.e. stems separated by an internal loop or bulge), L0 refers to length of the 5’-loop connecting H1 and H2;
2. *L1* & *L2*: Number of unpaired nucleotides at the terminal ends of H1 and H2, respectively. If H1 and H2 are directly connected, L1 = L2, both corresponding to the 3′-side loop length between the two stems;
3. *A(L0)*, *A(L1)*, *A(L2)*: Number of consecutive adenines within loops L0, L1, and L2, respectively (Nissen et al. 2001);
4. *L(H1)* & *L(H2)*: Lengths of stems H1 and H2, measured by the number of base pairs;
5. *N*: Number of stems located between H1 and H2 in the original n-way junction, reflecting their relative spatial separation;
6. Δ*G(H1)* & Δ*G(H2):* Free energy (ΔG) of stems H1 and H2, respectively.
7. Δ*G(H1-H2)*: Coaxial stacking free energy between H1 and H2, estimated from base stacking interactions at their junction interface (Ali et al. 2023).

To characterize the thermodynamic stability of each pseudo two-way junction, we computed three types of free energy features (*ΔG(H1)*, *ΔG(H2)*, and *ΔG(H1-H2)*) based on the RNA nearest-neighbor thermodynamic model developed by Turner’s group (Mathews et al. 2004). The free energy of each helix, *ΔG(H1)* and *ΔG(H2)*, was calculated using the Watson–Crick–Franklin (WCF) model, as a linear combination of initiation energy, AU-end penalties, symmetry correction, and base stacking contributions (Mathews et al. 2004):

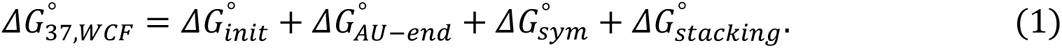

The coaxial stacking free energy between two adjacent helices, *ΔG(H₁–H₂)*, was estimated using the Turner nearest-neighbor parameters according to the following empirical function (Laing et al. 2012; Mathews et al. 2004; Mathews et al.1999):

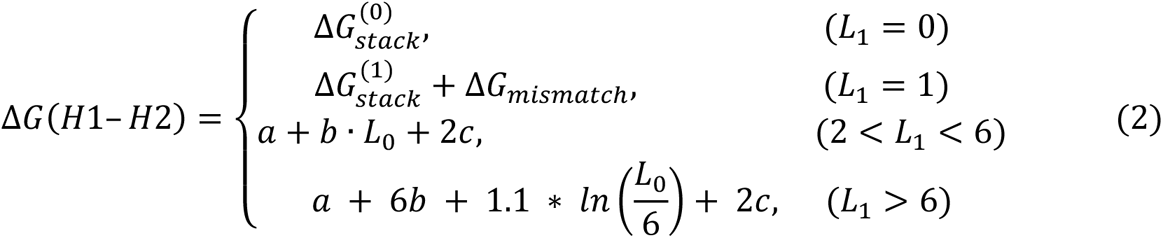

Here, 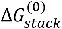 and 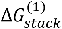 are coaxial stacking free energies with 0 and 1 unpaired nucleotide (i.e., *L0*) inserted between the two helices, respectively (Mathews et al. 2004; Tyagi and Mathews 2007). The term Δ*G*_*mismatch*_ in Eq. 2 represents the terminal mismatch penalty (2.1 kcal/mol). The empirical constants of *a*=9.3, *b*=−0.3, and *c*=0.9 were derived from Ref. (Mathews et al. 2004). Previous studies have shown that the likelihood of coaxial stacking decreases sharply with increasing loop length between stems (Lescoute and Westhof 2006; Tyagi and Mathews 2007). In particular, when *L0*≥2, stacking becomes thermodynamically less favorable. Accordingly, a penalty term is applied as formulated in Eq. 2 to reflect this reduced stacking propensity for longer loops.

### Training and Test Datasets

As shown in Fig. S9 in the Supplementary Materials, a total of 2,534 RNA-only structures were retrieved from the Protein Data Bank (PDB). To reduce sequence redundancy, we applied CD-HIT (Fu et al. 2012) with an 80% identity threshold, resulting in 681 representative RNA molecules (see Fig. S10). Since many complex RNAs (e.g., ribonucleases, group II introns, and ribosomal RNAs) contain multiple multi-way junctions, we used DSSR (Lu et al. 2015) to parse each 3D structure into distinct *n*-way junctions, disregarding tertiary interactions across different junctions.

To further reduce redundancy at the junction level, we clustered the resulting *n*-way junctions using CD-HIT with the same 80% sequence similarity threshold. This yielded a non-redundant set of RNA junctions for downstream analysis (Fig. S9). The final dataset includes 1642 two-way, 312 three-way, 231 four-way, 87 five-way, and 18 six-way or higher-order junctions. To balance the dataset, a random subset of 1457 two-way junctions was retained due to their overrepresentation. Each *n*-way junction was subsequently re-analyzed with DSSR to extract its secondary structure and annotated coaxial stacking interactions (Fig. S9). For model development, 85% of junctions from each n-way category were randomly assigned to the training set, and the remaining 15% formed Test Set I (Figs. S4 and S10). Notably, due to the limited number of six-way and higher-order RNA-only junctions in the non-redundant dataset, our test set included no RNA junctions beyond the five-way level. To evaluate the generalizability of gCoSRNA on higher-order structures, we additionally collected six-way and seven-way junctions from non–RNA-only structures in the original dataset. After removing any redundancy with the training set and within the test set itself, we obtained 45 six-way and 49 seven-way junctions. These were incorporated into Test Set I to assess the model’s performance on complex junction architectures beyond the training distribution. Finally, all training and test junctions were decomposed into pseudo two-way junctions and processed through the same feature extraction pipeline, yielding the final dataset used for model training and evaluation.

To further evaluate model generalizability, we assembled an independent Test Set II from RNA-Puzzles and CASP 15/16 targets (Bu et al. 2025; Das et al. 2023; Kretsch et al. 2025). After removing RNAs sharing more than 80% sequence identity with the training set, this dataset consisted of 80 two-way, 9 three-way, and 29 four-way junctions (Figs. S4C & D). These junctions were similarly decomposed and featurized using the same protocol.

### Model Training and Inference Strategy

Using the constructed dataset, we trained a binary Random Forest (RF) classifier to predict whether two stems within a pseudo two-way junction are coaxially stacked. Given the high-dimensional and limited-sample nature of RNA coaxial stacking data, we adopted a Bayesian optimization strategy to tune model hyperparameters efficiently. Six key hyperparameters were optimized: the number of trees, maximum tree depth, minimum samples required to split an internal node, minimum samples at a leaf node, feature sampling ratio, and the class weight assigned to positive samples. A Gaussian process model, coupled with 10-fold cross-validation, was employed to explore the hyperparameter space. The optimization procedure began with five random initializations, followed by ten iterations of guided search. The final optimized configuration was as follows: 562 trees, a maximum depth of 24, minimum split size of 18, minimum leaf size of 1, feature sampling ratio of 0.1, and a positive class weight of 1.09.

During inference, each n-way junction is decomposed into all possible adjacent pseudo two-way junctions. For each pair, the trained RF model outputs a binary stacking label and an associated probability *p*. The complete coaxial stacking configuration is then reconstructed as follows (Fig. 1A): (1) stem pairs with *p* ≥ *τ* (a user-defined threshold, e.g., 0.42 used in this work; see Fig. S11) are considered coaxially stacked; (2) if a single stem is predicted to stack with multiple partners, only the pair with the highest *p* is retained; (3) the retained stem pairs are integrated to produce the final coaxial stacking topology of the *n*-way junction.

### Re-implementation of RNA Junction-Explorer for Benchmarking

RNA Junction-Explorer is a Random Forest–based method originally developed to predict coaxial stacking configurations in RNA junctions using features derived from secondary structure (Laing et al. 2012). It remains the only publicly reported tool specifically designed for this purpose. The method extracts junction-level features (e.g., 15, 18, and 10 features for three-, four-, and high-order-junctions), such as loop length, nucleotide composition, and helix-end stacking energies, and trains separate classifiers for junctions of different orders (e.g., three-way, and four-way) (Laing et al. 2012). However, the official web server (https://nature.njit.edu/biosoft/Junction-Explorer/) is no longer functional and fails to return valid results, likely due to discontinued maintenance. To facilitate a direct comparison with our method gCoSRNA, we re-implemented the RNA Junction-Explorer pipeline according to the algorithmic details described in its original publication (Laing et al. 2012).

We first replicated model training using the original feature sets and training data (e.g.,110 three-way junctions), along with the hyperparameters reported in the original RNA Junction-Explorer publication (Laing et al. 2012). To enable a more rigorous comparison, we then retrained the model, referred to as Junction-Explorer-n, using our newly curated and non-redundant RNA junction dataset, while keeping the original feature definitions unchanged. This updated dataset increases the number of training samples by 138% for three-way junctions and 194% for four-way junctions, respectively, relative to the original corpus. For retraining, we applied Bayesian optimization with 10-fold cross-validation to re-tune model hyperparameters for improved generalizability.

### Evaluation Metrics

To comprehensively assess the performance of gCoSRNA across different junction orders and classification settings, we employed a combination of standard and task-specific metrics (see Supplementary Material for details). For the binary classification task of predicting coaxial stacking between two stems in (pseudo) two-way junctions, we reported accuracy, F1-score, and area under the ROC curve (AUC), all computed from the standard confusion matrix. In the multi-class setting for three-way and four-way junctions, where the number of possible stacking topologies increases to four and seven, respectively (Fig. S3). In addition, we included the full confusion matrices to highlight specific patterns of misclassification.

For junctions with five or more stems, the combinatorial complexity of full-topology classification becomes prohibitive. Following the convention used in RNA Junction-Explorer (Laing et al. 2012), we instead focused on the accuracy of local pairwise stacking decisions between adjacent stem pairs. Specifically, we defined the pairwise accuracy (*PA*) as:

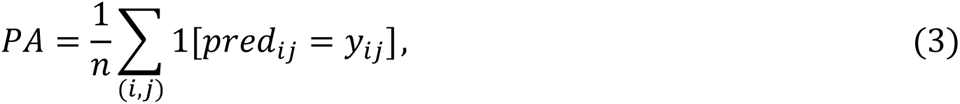

where *n* is the number of cyclically adjacent stem pairs (*H_i_*, *H_i+1_*) in an *n*-way junction, *y_ij_* denotes the ground truth (stacked/unstacked), and *pred_ij_* is the model prediction (see Fig. S12). This metric captures the fidelity of local stacking predictions while remaining tractable for higher-order junctions.

## SUPPLEMENTAL MATERIAL

Supplemental material is available for this article.

## ACKNOWLEDGMENTS

We are grateful to Profs. Zhi-Jie Tan (Wuhan University), and Yaoqi Zhou (Shenzhen Bay Laboratory) for valuable discussions, and we would like to acknowledge computing resources from the Super Computing Center of Wuhan Textile University. This work was supported by the National Natural Science Foundation of China [grant numbers 12371500, 12271416, and 1220522], the Department of Education of Hubei Province [grant number Q20221705], and the Special Research Fund of Wuhan Textile University.

